# Dynamic Domain Specificity In Human Ventral Temporal Cortex

**DOI:** 10.1101/2020.11.11.378877

**Authors:** Brett B. Bankson, Matthew J. Boring, R. Mark Richardson, Avniel Singh Ghuman

## Abstract

An enduring neuroscientific debate concerns the extent to which neural representation is restricted to neural populations specialized for particular domains of perceptual input, or distributed outside of highly selective populations as well. A critical level for this debate is the neural representation of the identity of individual images, such as individual-level face or written word recognition. Here, intracranial recordings throughout ventral temporal cortex across 17 human subjects were used to assess the spatiotemporal dynamics of individual word and face processing within and outside regions strongly selective for these categories of visual information. Individual faces and words were first discriminable primarily only in strongly selective areas, beginning at about 150 milliseconds after word or face onset, and then discriminable *both* within and outside strongly selective areas approximately 170 milliseconds later. Regions of ventral temporal cortex that were and were not strongly selective both contributed non-redundant information to the discrimination of individual images. These results can reconcile previous results endorsing disparate poles of the domain specificity debate by highlighting the temporally segregated contributions of different functionally defined cortical areas to individual level representations. This work supports a dynamic model of neural representation characterized by successive domain-specific and distributed processing stages.

## INTRODUCTION

A key debate regarding the architecture of the cortex concerns the extent to which diagnostic aspects of stimuli are processed within domain-specific cortical populations (Kanwisher et al., 1997; Martin, 2007; Fodor, 1983), or if processing is also distributed outside of highly selective neural populations (Haxby et al., 2001; Op de Beeck, 2008). On one hand, an extensive body of primate single unit recordings (Perrett et al., 1984; Tsao et al., 2006), human neuroimaging (Kanwisher et al., 1997; Puce et al., 1996), stimulation (Puce et al., 1999; Hirshorn et al., 2016; Afraz et al., 2006; Pitcher et al., 2007; Schalk et al., 2017), and lesion (Farah et al., 1995; Hirshorn et al., 2016; Gaillard et al., 2006) data suggests that perception is causally related to the activity within systems of cortical populations that respond selectively to preferred stimulus categories. Conversely, the distributed representation hypothesis is supported by evidence from both neuroimaging and single unit recordings that shows reliable face differentiation in weakly or non-face-selective populations (Haxby et al, 2001; Bell et al., 2011) and differentiation of non-face categories within face selective populations (Kiani et al., 2007; Cukur et al., 2013; Hanson & Schmidt, 2011).

Across these hypotheses, a central point of debate concerns the role of activity evoked by stimuli outside of highly selective parts of VTC (e.g. face-related activity outside of highly face selective populations) and activity evoked by “other” stimuli inside parts of VTC selective for particular categories of stimuli (e.g. non-face activity in face selective populations). A critical tension between the aforementioned hypotheses is whether individual-level discrimination (e.g. recognizing which face or word a person is viewing) can be found outside of putative category-selective regions of VTC (Spiridon & Kanwisher, 2002; Nestor et al., 2011). Because individual-level perception, but not category-level discrimination, is compromised in various agnosias (Damasio et al., 1982), addressing the debate between domain specific and distributed models of processing requires the comparison of individual-level representations inside and outside of parts of VTC that are highly selective at the category level (Spiridon & Kanwisher, 2002).

To test for the presence of individual-level representations across time in and out of highly selective regions, the dynamics of face individuation was examined with intracranial electroencephalography (iEEG) in 14 patients with pharmacologically intractable epilepsy. To ensure that face individuation was based on face identity level and not the visual image level, 15 different images of each of 14 different identities were used across 5 expressions (anger, sadness, fear, happy, neutral) and 3 gaze directions (left, straight, right). The dynamics of word individuation was examined in 5 patients (2 overlapping, 17 total patients in the study). Face and word stimuli were chosen because 1) they comprise domains of visual stimuli for which human adults demonstrate strong expertise in exemplar-level discrimination, but 2) faces have a putative genetic basis that can be seen in our evolutionary ancestors and that infants are predisposed to orient to (Powell et al., 2018), and word expertise must be acquired during development. Thus, if similar findings are seen for both faces and words, it supports a general principle of organization for both learned and putatively partially innate information processing.

Above chance classification of individual faces and words was seen in both high face and word selective regions (HFS and HWS) and not-highly face and word selective regions (NFS and NWS), but significant decoding emerged approximately 170 ms earlier in HFS and HWS compared to NFS and NWS regions. These results suggest a dynamic model of domain specificity in VTC in which processing is first restricted to highly selective parts of VTC and then is processed a non-redundant, though also partially similar, manner inside and outside of highly selective regions.

## RESULTS

### Spatiotemporal dynamics of individuation

Significant face and word individuation were present in and out of HFS and HWS regions (Figure 2), as measured with elastic net regularized logistic regression. Using the first method of onset calculation (see methods under Statistical Analysis), the onset of face individuation occurred 190 ms earlier in HFS regions relative to NFS regions (*t*(13)= 3.05, *p* = 0.009) and peaked 200 ms earlier (*t*(13) = 2.73, *p* = 0.017), with a higher peak in HFS than NFS regions (*t*(13) = 2.68, *p* = 0.019). Notably, the difference in the magnitude of the HFS and NFS response is independent of the difference in peak times, though onset times can be affected by magnitude differences. Using two other methods of onset calculation that are robust to differences in magnitude (Schrouff et al., 2020), above chance face individuation occurred significantly earlier inside (160 ms, 210 ms) than outside (250 ms, 325 ms) HFS regions (*t*(13) = 3.6, *p* = 0.003; *t*(13) = 3.03, *p* = 0.0096).

**Figure 1.**
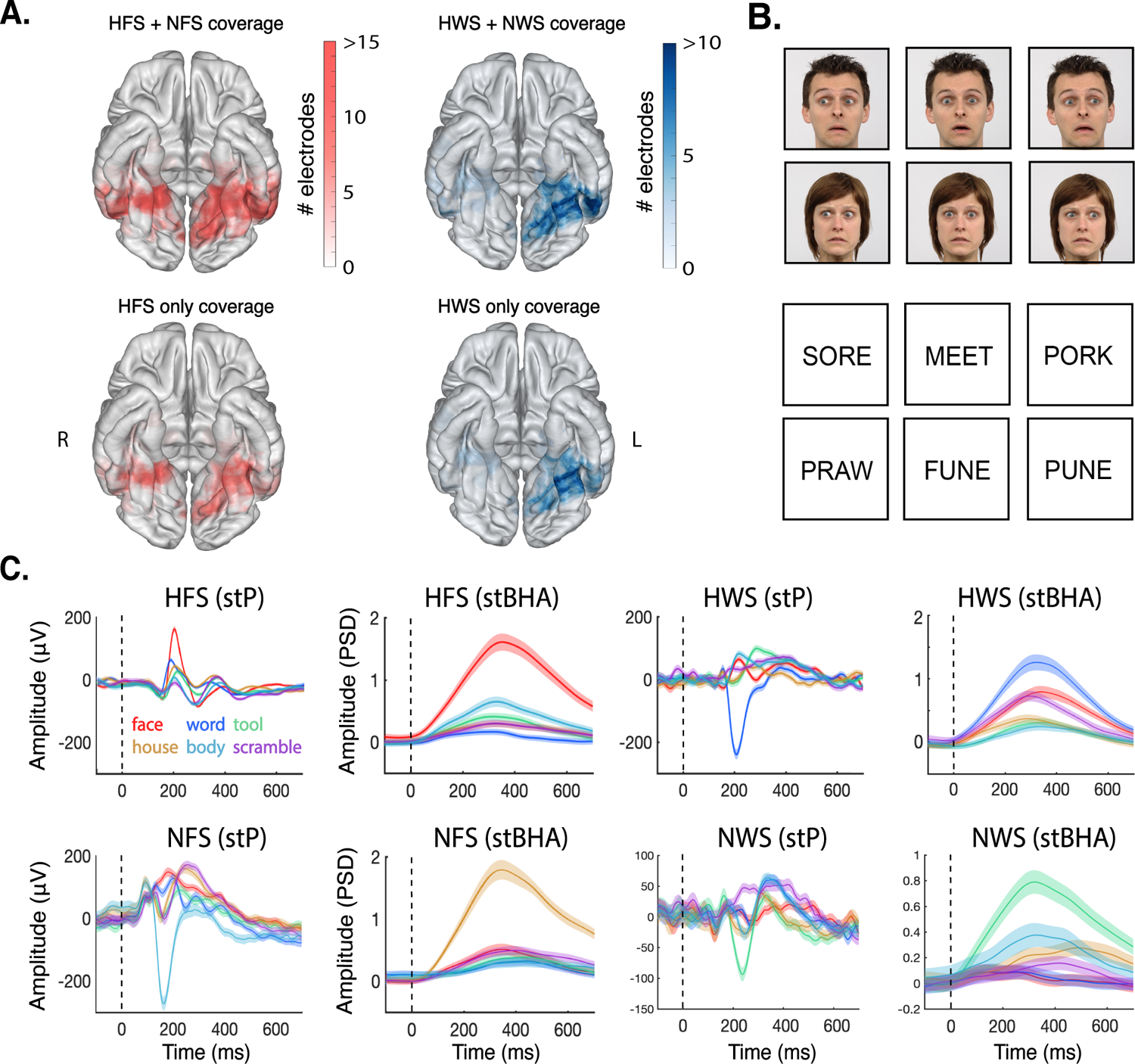
Ventral temporal electrode contact heatmaps (489 total electrode contacts), stimulus examples, and electrophysiological traces. **A)** 426 total contacts from 14 subjects who completed the gender discrimination task were divided into 171 HFS and 255 NFS contacts. Total HFS + NFS coverage throughout ventral temporal cortex is depicted in the top left heatmap. Total HFS coverage is depicted in the bottom left heatmap. 174 total contacts from 5 subjects who completed the word one-back task were divided into 113 HWS and 61 NWS contacts. Total HWS + NWS coverage throughout ventral temporal cortex is depicted in the top right heatmap. Total HFS coverage is depicted in the bottom right heatmap. Electrode contact locations depicted here include both subdural electrode strips on the surface of the cortex and stereotaxically implanted depth electrodes, projected to the nearest surface vertex. **B)** Example stimuli from gender discrimination task demonstrating 2 male and female identities with 3 gaze directions and a surprised expression. Example stimuli from word one-back task depicting four-letter real and matched pseudowords. **C)** Averaged single trial potentials (measured in microvolts) and single trial high broadband activity (measured with z-scored power spectrum density values) from single electrodes from high face selectivity, high word selectivity, non-high face selectivity, and non-high word selectivity populations in response to 6 visual categories during localizer task (note that for subjects that performed both tasks, HFS electrode contacts are included in the NWS group and HWS contacts are included in the NFS group). Dashed black line shows stimulus onset. See Boring et al. (2021) for a full examination of the localization and category-level neurodynamics of face and word selective electrodes in iEEG in a superset of subjects that include the ones reported in this work on the dynamics of individual level face and word processing.

**Figure 2.**
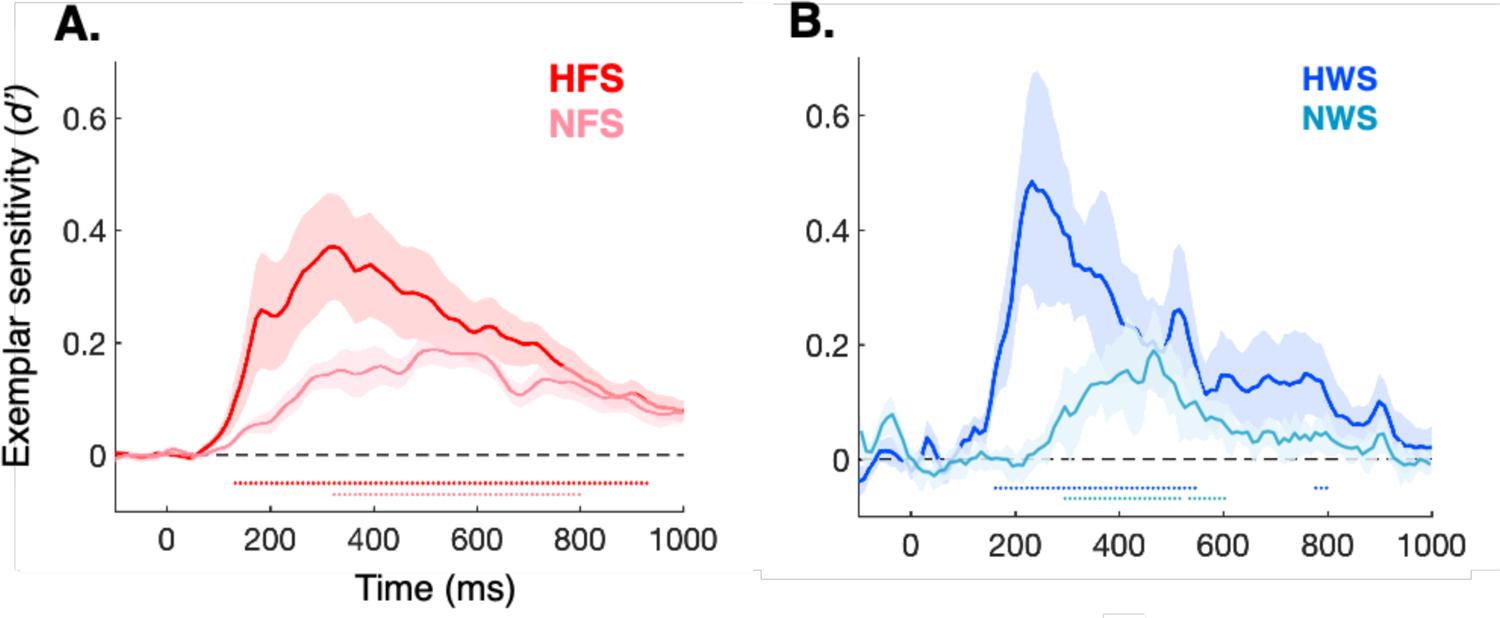
Time course of the sensitivity index (*d’*) for individuation of faces and words. **A)** HFS contacts (dark red) demonstrated significant face individuation from 130 ms to 940 ms after stimulus onset, with peak *d’* = 0.37 at 320 ms (*p* < .05, FDR corrected for multiple dependent temporal comparisons). NFS contacts (light red) demonstrated significant face individuation from 320 ms to 800 ms after stimulus onset, with peak *d’* = 0.17 at 520 ms (*p* < .05, FDR corrected). HFS individuation onset emerged significantly earlier than NFS individuation (190 ms average difference between HFS and NFS onset, *p* = .009, *t(13)* = 3.05). Individually, 11 of the 14 patients demonstrated an earlier onset of significant individuation in HFS than NFS contacts. **B)** HWS contacts (dark blue) demonstrated significant word individuation from 160 ms to 535 ms after stimulus onset, with peak *d’* = 0.48 at 235 ms (*p* < .05, FDR corrected for dependent tests). NWS contacts (light blue) demonstrated significant word individuation from 285 ms to 605 ms after stimulus onset, with peak *d’* = 0.18 at 470 ms (*p* < .05, FDR corrected). Word individuation emerged significantly earlier in HWS compared to NWS regions (145 ms average difference between HWS and NWS onset, *p* = .036, *t(4)* = 3.1). Individually, all 5 patients demonstrated an earlier onset of word individuation in HWS compared to NWS contacts. Shaded bars illustrate standard error of the mean across subjects at each time point.

Word individuation began 145 ms earlier in HWS regions relative to NWS regions (*t*(4) = 3.1, *p* = 0.036) and peaked 250 ms earlier (*t*(4) = 3.61, *p* = 0.022), with a higher peak in HWS than NWS regions (*t*(4) = 2.802, *p* = 0.048). Using the two other methods of onset calculation that are more robust to differences in magnitude (Schrouff et al., 2020), above chance face individuation occurred earlier inside (150 ms, 190 ms) than outside (285 ms, 405 ms) HFS regions (*t*(4) = 1.77, *p* = 0.15; *t*(4) = 4.31, *p* = 0.01).

HFS and HWS regions maintained significant sensitivity to individual face and word information respectively throughout visual processing (from 130-940 ms and 160-535 ms respectively, *p*<0.05 FDR corrected), suggesting that these regions contribute to both early and late visual processing (before and after 300 ms). NFS and NWS reached significance only later (from 320-800 ms and 285 - 605 ms respectively, *p* < 0.05 FDR corrected), suggesting that these regions contribute to late visual processing. For both faces and words, the finding of earlier individuation in high selectivity regions relative to non-highly selectivity regions was robust across a range of criteria for defining “highly” and “non-highly” selective (Figure 3). The robustness of the result demonstrates that illustrating that the differences in timing were not due to choosing an arbitrary threshold for “high” selectivity.

**Figure 3.**
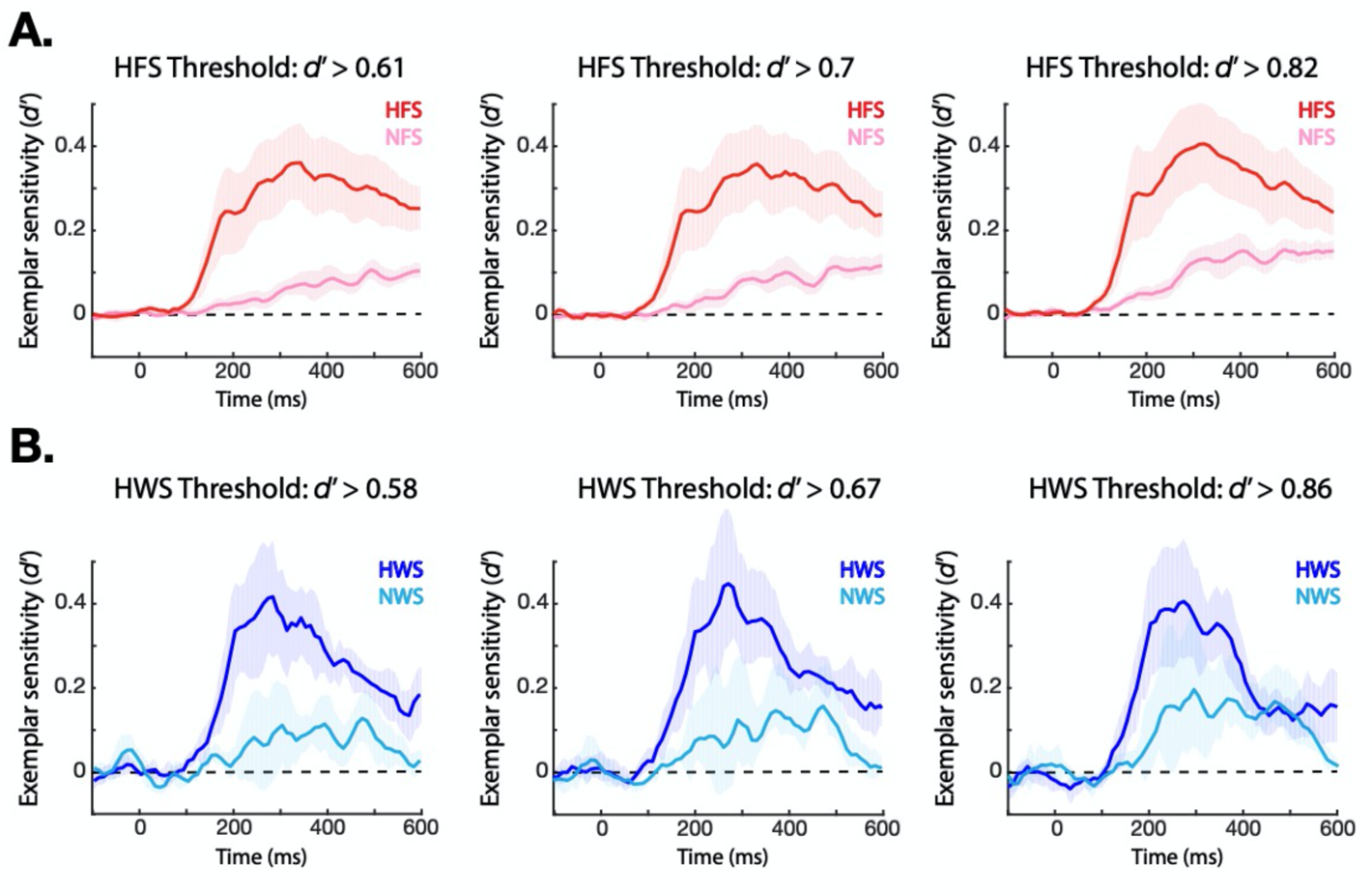
Time course of the sensitivity index (*d’*) for individuation of faces and words at graded thresholds. Individual face and word classification (as in Figure 2) was repeated with multiple “high” selectivity thresholds. These thresholds were defined by separating all contacts into partitions corresponding to the one third, one half, and two-thirds levels of d’ values as measured with the category localizer. **A)** Time course of face individuation at thresholds of *d’* = 0.61 (NFS = bottom 1/3 of contacts, HFS = top 2/3 of contacts), *d’* = 0.7 (NFS = bottom ½ of contacts, HFS = top ½ of contacts), and *d’* = 0.82 (NFS = bottom 2/3 of contacts, HFS = top 1/3 of contacts). **B)** Time course of word individuation at thresholds of *d’* = 0.58 (NWS = bottom 1/3 of contacts, HWS = top 2/3 of contacts), *d’* = 0.67 (NWS = bottom ½ of contacts, HWS = top ½ of contacts), and *d’* = 0.86 (NWS = bottom 2/3 of contacts, HWS = top 1/3 of contacts).

Electrodes were placed based on the clinical needs of the patients and not necessarily optimally placed for sensitivity to visual information, thus relative effect sizes are likely more relevant than absolute effect sizes. Peak effect sizes in NFS and NWS regions were relatively small, but nonetheless more than 1/3 that of the peak effect sizes in HFS and HWS regions. This suggests that activity in NFS and NWS regions contributed meaningfully to the overall representation of individual faces and words, albeit less than HFS and HWS regions. Every patient had recordings from both highly and non-highly category-selective areas.

To address potential concerns of signal bleed as the source of face individuation signals in non-highly selective populations, we used multivariate regression to remove all of the high selectivity channels’ activity from the non-highly selective channels and examined whether the residual signals showed above chance classification. In both NFS and NWS regions, the residual activity showed significant individuation during the stimulus presentation period (*p* < .05 FDR corrected) after regressing out the multivariate signal from HFS and HWS channels.

The regression analysis above demonstrates that NFS and NWS contain at least some diagnostic face and word information that is not redundant to the information in HFS and HWS regions. The complimentary question is whether there is some shared information between high and non-highly selective regions as well. To address this question, we used RSA to show that HFS and NFS populations share significant overlap in face individuation structure (*p* < .01, *t*(13) > 3.17, and HWS and NWS populations share significant overlap in word individuation structure (*p* < .01, *t*(4) > 4.68). Thus, non-highly and highly selective regions have both some unshared information (based on significant classification in non-highly selective regions after regressing out the activity from high selectivity regions) and some shared information (based on significant correlation in the RSA analysis).

### Relative contribution of highly and non-highly selective regions to individuation

The previous results demonstrate that individuation emerges earlier inside highly selective regions than outside these regions, but leaves the relative contribution of activity in highly and non-highly selective regions to the overall individual-level representation unclear. Specifically, two important questions are outstanding: 1) What is the balance of information between non-highly selective regions and highly selective regions? 2) Outside of highly selective regions, to what extent is discriminant information present in regions that are selective to other categories or regions that show no measured category selectivity, e.g. do word-selective contacts (or body, or house, etc. selective contacts) contribute diagnostic information to face individuation?

Regarding the first question, the multivariate regression results discussed above show that NFS and NWS regions contain signals discriminant for faces and words beyond what is present in HFS and HWS regions, but does not assess the relative information in each. To address this question, sparse classification using L1-regularization and identical parameters to earlier elastic net procedure except regularization parameter (λ) was performed over all ventral temporal contacts to identify the electrode contacts that provided information for face or word individuation. If activity between any set of contacts is highly correlated, L1-regularization should force all contacts in that set to have zero weight, except the one with the largest amount of discriminating information. Thus, the balance of non-highly and highly selective electrodes that survive L1-regularization provides an estimate of how much each population of electrodes contribute to the overall information about individual faces and words in VTC as a whole. Note that choice of regularization method (elastic net vs. L1) does not alter the pattern of reported results above (supplemental Figure 1). To address the second question, the above analysis was extended by decomposing the non-highly selective contacts into “other category-selective” (OCS) and “not category-selective” (NCS) populations. This was done by identifying the NFS and NWS contacts that showed high selectivity for any of the other 5 categories in the localizer and those that did not.

For both face and word individuation tasks, the analysis showed that proportions of both HFS/HWS and NFS/NWS electrode populations contribute diagnostic information (Figure 4A), though highly selective regions may contribute more than non-high selectivity ones. Second, decomposing the NFS contacts showed that in the face individuation task, regions highly selective for other categories contribute diagnostic information to overall individuation as well as those that demonstrate non-high selectivity for all categories (Figure 4B). Specifically, higher proportions of OCS than NCS electrode contacts survive penalization and contribute diagnostic information to exemplar classification using L1 regularization. These findings demonstrate that in the later time period, meaningful information that contributed to above chance individuation is present outside of category-selective areas, distributed even to areas that demonstrate selectivity for a different visual object category.

**Figure 4.**
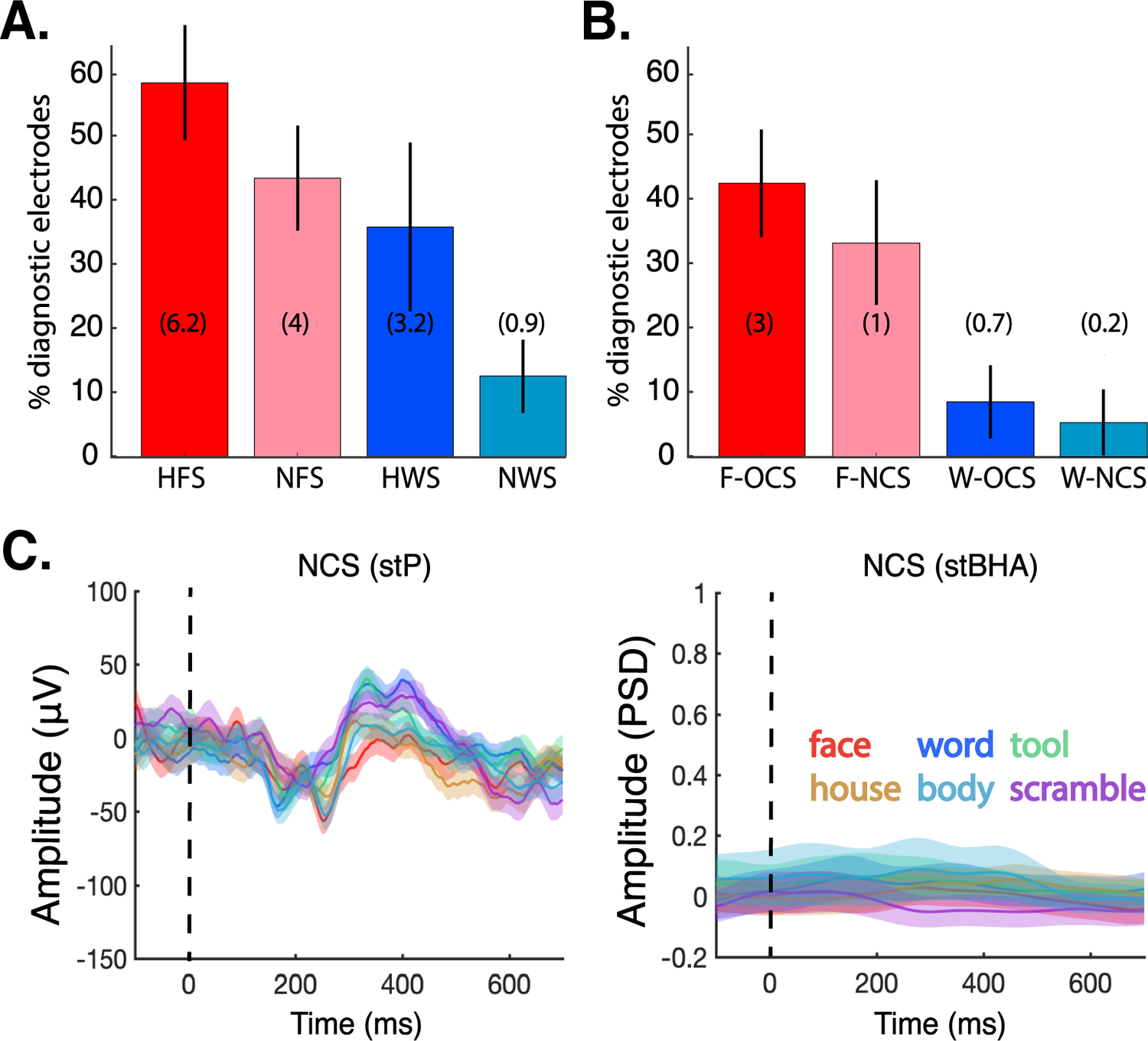
**A)** Percent diagnostic electrode contacts inside and outside high category-selectivity populations measured with L1-regularized logistic regression. 58.6% of HFS contacts from 14 patients (average of 6.2 contacts per subject, SEM = 9%) were assigned non-zero weights. 42.4% of NFS contacts (average of 4 contacts, SEM = 8.9%) were assigned non-zero weights on average. 35.2% of HWS contacts from 5 patients (average of 3.2 contacts, SEM = 18%) on average were assigned non-zero weights. 11.4% of NWS contacts (average of 0.9 contacts, SEM = 6.2%) were assigned non-zero weights. **B)** Within non-high face selectivity electrode contacts that demonstrated selectivity for a different object category (F-OCS, other category-selective), 42.5% of the contacts (average of 3 contacts per montage, SEM = 8.7% of contacts) were assigned non-zero weights. Within non-high face selectivity contacts that demonstrated no selectivity for any object categories (F-NCS, not category-selective), 33% of the contacts (average of 1 contact, SEM = 9.6%) on average were assigned non-zero weights. Within non-high word selectivity electrode contacts that demonstrated selectivity for a different object category (W-OCS), 9.4% of the contacts (average of 0.7 contacts, SEM = 8.1%) were assigned non-zero weights. Within non-high word selectivity contacts that demonstrated non-significant selectivity for any other object categories (W-NCS), 5% of the contacts (average of 0.2 contacts, SEM 5%) on average were assigned non-zero weights. **C)** Example single trial potential (microvolts) and single trial high broadband activity (z-scored power spectrum density values) traces of one NCS electrode contact in response to all 6 categories of localizer task stimuli. The dashed black line indicates stimulus onset. Example single trial potential and single trial high broadband activity traces of one OCS electrode contact (showing tool selectivity) in response to all 6 categories of localizer task stimuli can be seen in the NWS panels of figure 1C.

## Discussion

The presence of individual-level information in and out of highly category-selective electrode contacts at different latencies suggests a “dynamic domain specificity” model of visual processing. Specifically, information from a given visual category is first processed primarily in strongly category-selective cortical populations followed by widespread processing that includes both populations that are strongly and weakly selective for that stimulus category (Shehzad & McCarthy, 2018). The cascade of neural activity during visual perception is characterized by an early, potentially obligatory, stage of processing in strongly category-selective regions that may guide and gate information for further processing. Previous studies suggest that this early stage represents a coarse pass of processing only allowing for differentiation of relatively distinct images (Hirshorn et al., 2016; Ghuman et al., 2014; Hegdé, 2008). Approximately 150-200 ms later, information then flows to visual processing populations outside of strongly category-selective populations as well, including into cortical populations that are selective for other visual categories, either through lateral and recurrent connectivity or through top-down feedback. Non-highly selective regions contribute unique information to the overall individual-level representation, though both these and high selectivity regions also exhibit partial representational overlap. Future studies, perhaps requiring single unit recordings (Chang & Tsao, 2017), will be required to determine the precise nature of the similarities and differences in the representational structure for faces and words in non-highly versus highly selective regions. The extra processing capacity from non-highly selective regions may help support later visual processing (Hirshorn et al, 2016; Ghuman et al., 2014; Li et al., 2019) that could contribute to determining subtle distinctions between individual category members or assist with later processes coincident to the time when activity from non-highly selective regions begin to show significant individuation, such as viewpoint or position generalization (Freiwald et al., 2010; Quian Quiroga, 2012; Mormann et al., 2008, Quian Quiroga, 2005; Tang et al., 2014).

The proposed dynamic domain specificity hypothesis may reconcile apparent contradictions between findings that have been used to support domain-specific and distributed models of visual perception. The profound and frank disturbances to the perception of stimuli from particular categories seen in the presence of lesions or disruptions to highly category-selective regions (Puce et al., 1999; Parvizi et al., 2012; Afraz et al., 2006; Farah et al., 1995; Schalk et al., 2017; Rangarajan et al., 2014) may emerge due to the perturbation of early and potentially obligatory activity of these areas during visual processing. The perceptual relevance of later activity in non-highly selective regions is supported by the current evidence that these regions contribute some unique information to face and word individuation (Figure 4 and significant classification in non-highly selective regions after regressing out activity from high selectivity regions). The time of peak individuation in non-highly selective regions occurs when significant individuation is still present in high selectivity regions and is near the time when key higher-level visual processes such as viewpoint generalization (Freiwald et al., 2010) and semantic processing (Clarke et al., 2015) occur. Additionally, single units in the medial temporal lobes show selectivity for individual faces in a similar later time period and it has been suggested that this time period is critical for linking perception and memory (Quian Quiroga, 2012; Mormann et al., 2008, Quian Quiroga, 2005). Furthermore, this time window is substantially earlier than behavioral reaction times for comparable individual-level face and word recognition tasks (Haxby et al., 1999; Seidenberg & McClelland, 1989). The later information processing in non-high selectivity regions would also help explain why category discriminant information is sometimes seen outside of category-selective regions in low temporal resolution measures such as fMRI (Haxby et al., 2001; Ghuman & Martin, 2019). As such, non-highly selective regions may play a role in some aspects of individuation, even if that role is later and more supportive than the central role of strongly selective regions.

A recent study showed that electrical stimulation to NFS electrode contacts does not cause frank distortions of face perception (Rangarajan et al., 2014) and stimulation to NWS electrode contacts does not cause frank distortions of reading (Hirshorn et al., 2016), though these studies were not sensitive to the subtle aspects of perception that may be caused by disrupting areas that play a supportive role in processing. Causal manipulations of activity in non-highly selective regions, particularly ones that were precisely timed relative to stimulus onset, coupled with measures of subtle aspects of perception in the future would be useful to determine what role non-highly selective regions may play in individuation. One alternative explanation of later discrimination in non-highly selective regions that would support a non-causal role in perception is that it could reflect a backpropagating learning signal (Rumelhart et al., 1986) rather than perceptual processing per se.

While the results here are consistent with the primarily low temporal resolution data that have been used to support both domain specific and distributed models of VTC organization, they also help address theoretical aspects of the debate between the models. Specifically, in distributed models the difference between strongly and less selective parts of VTC is a difference in the degree to which each contributes to perception of stimuli from a particular category, but these contributions should happen at the same processing stage. These models would predict that highly and non-highly selective regions should each have similar timecourses of processing, varying mostly in how much each contributes to the representation for a particular stimulus class. The result that individual-level representations in highly selective regions onset and peak 145 - 250 ms earlier than in non-highly selective regions presents a challenge to current instantiations of distributed models. These differences survive across a range of criteria for selectivity (Figure 3), suggesting there is a qualitative, not graded, difference in the role that highly selective regions play for processing stimuli that those regions are selective for relative to non-highly selective regions. Thus, distributed models would need to be modified to accommodate relationships between selectivity and latency of information processing. One possibility that our results cannot exclude is that there is a continuous relationship between selectivity and timing of individual-level information rather than a bivariate one. If the relationship was continuous, it would suggest that the regions with the strongest selectivity contribute throughout perceptual processing, moderate selectivity regions contribute from a middle stage through the end, and weakly selective regions only for the longest latency processes.

In the strongest versions of domain specificity models, there is no role for parts of VTC that are not highly selective for a particular category of image in perceptual processing for that stimulus type. However, the results here suggest that these non-highly selective regions do contribute to later visual processing. The dynamic domain specificity hypothesis outlined above is an attempt to modify traditional models of domain specificity by positing a supportive role for non-highly selective regions; they may support later processes and provide supplementary computational resources may be particularly useful in aiding more difficult perceptual processes.

The dynamic pattern of results was seen for both faces, with circuitry that putatively arises from evolutionary and genetic origins, and words, where reading skill must be acquired fully through experience, suggesting that dynamic domain specificity may be a general principle of cortical organization. One caveat is that words were not varied with regards to visual appearance. Thus, word individuation results may reflect discrimination of visual features and our results cannot rule out that dynamic domain specificity may not apply to words per se and may only apply to word-like shapes. Nonetheless, the results with words still provide support for the generalizability of dynamic domain specificity as it shows this principle governs an additional well-learned category other than faces.

Taken together, these results may reconcile the tension between domain-specific versus distributed models of visual object processing by providing evidence that domain-specific and distributed processing emerge dynamically at different times during the course of visual perception.

## MATERIALS AND METHODS

### Subjects

Experimental protocols were approved by the Institutional Review Board of the University of Pittsburgh and written informed consent was obtained from all subjects. 17 patients (8 female) undergoing surgical treatment for medicine-resistant epilepsy volunteered to participate in this experiment. Patients had previously undergone surgical placement of intracranial surface / grid and/or stereotactic electroencephalography depth electrodes (collectively referred to as iEEG here) as standard care for clinical monitoring during seizure onset zone localization. 13 of the 17 patients exclusively had stereotaxic depth electrode implantations, and the remaining 4 patients had a combination of grid / strip surface electrodes on cortical regions and depth electrode implantations in subcortical structures. For stereotaxic depth electrodes, each adaptor contained 32 electrode contacts with 1 common reference and 3 ground contacts that were used to normalize signal in each set of 32 contacts. For grid and strip surface electrodes, the first 2 contacts for each grid (differing numbers of contacts depending on custom dimensions) were used to reference and ground the grid signal.

Only patients with grid electrodes had craniotomies performed over the target cortical tissue. Depth electrodes were produced by Ad-Tech Medical and PMT Corporation and the electrode contacts were 0.86 and 0.8 mm in diameter, respectively. Grid electrodes were produced by PMT Corporation and the electrode contacts were 4 mm in diameter. Because depth electrode contacts are cylindrical, the surface area of the recording site was similar across grid and strip electrode contacts. Post-operative MRIs were performed for patients with depth electrodes, but standard clinical procedure follows a pre-operative MRI and post-operative CT for patients with grids because grids electrodes are difficult to visualize using MRI. All patients underwent standard post-operative clinical procedures for recovery and experiments were run at least 36-48 hours after surgery to ensure adequate post-operative recovery. Recordings all took place in the UPMC Presbyterian Epilepsy Monitoring Unit in Pittsburgh, PA. Local field potentials were recorded via a GrapeVine Neural Interface (Ripple, LLC) sampling at 1 kHz. The amplification system used was a Natus Xltek 128-channel Brain Monitor EEG Amplifier.

The ages of subjects ranged from 20 to 64 years (mean = 39.1, SD = 14.6). None of the subjects showed any ictal events on any electrodes during experimental recording nor did they have epileptic activity on the electrodes used in this study at any time. All patients completed a localizer session, 14 patients completed experiment 1, and 5 patients (2 overlap) completed experiment 2.

### Experimental Design: Stimuli

In the localizer session, images of 6 categories (bodies (50% male), faces (50% male), words, hammers, houses and phase scrambled faces) were presented in a 1-back exact image repeat detection task. Specific examples of these stimuli are outlined in Figure 2 of Ghuman et al. (2014). Phase scrambled images were created in Matlab by taking the two-dimensional spatial Fourier spectrum of the image, extracting the phase, adding random phases, recombining the phase and amplitude, and taking the inverse two-dimensional spatial Fourier spectrum. Each image category was presented 80 times, yielding a total of 480 image presentations. Each image was presented for 900 ms, with a 900 ms inter-stimulus interval in pseudorandom order and repeated once in each session.

For experiment 1, frontal views of 14 different face identities were drawn from the Radboud Faces Database. 15 images of each identity were presented, with five expressions (anger, sadness, fear, happy, neutral) and three gaze directions (left, right, forward). Each unique image was presented four times, yielding a total of 60 presentations per identity and 840 face image presentations. For experiment 2, 36 different character strings corresponding to real words of 3-4 characters, pseudo-words of 4-5 characters (pronounceable letter strings that do not form real words, such as “lerm”), and false font words of 5 characters were presented 30 times each.

Pseudowords were selected to have similar mean bigram and trigram frequency as real words (measured using the English Lexicon Project). Because three-letter words did not have any corresponding pseudo-word stimuli, only trials from the 16 unique four-letter real and pseudo-word stimuli were considered further for data analysis. Thus, 480 trials of pronounceable orthographic stimuli were ultimately included in further analyses. All stimuli for the three experimental sessions were presented on an LCD computer screen placed ∼1 meter from subjects’ heads. Stimulus examples are shown in Figure 1B.

### Experimental Design: Paradigms

In all experimental sessions, each image was presented for 900 ms with 900 ms inter-trial interval during which a fixation cross was presented at the center of the screen (∼10° × 10° of visual angle for the localizer session and experiment 1, ∼6° × 6° visual angle for experiment 2). For the localizer session, images were repeated 20% of the time at random. Subjects were instructed to press a button on a button box when an image was repeated (1-back). Only the first presentations of repeated images were used in the analysis.

In experiment 1, subjects completed a gender discrimination task, reporting whether the presented face was male or female via button press on a button box. Each subject completed one or two sessions of the task. All three paradigms were coded in MATLAB (version 2007, Mathworks, Natick, MA) using Psychtoolbox (Brainard, 1997) and custom written code.

In experiment 2, subjects completed a one-back task, reporting whether the presented word (real or pseudo-word comparisons) was the same as the previous image via button press on a button box. Each subject completed one or two sessions of the task. All three paradigms were coded in MATLAB using Psychtoolbox and custom written code.

### Data preprocessing

Preprocessing followed the general steps of signal acquisition, trial segmentation from signal epochs, band-pass filtering to yield single trial potentials, and power spectrum density estimation to yield single trial broadband high-frequency activity. Electrophysiological activity was recorded at 1000 Hz using iEEG electrodes. These data were then epoched from −500 to 1500 ms trials around stimulus onset. Single-trial potentials were generated by band-pass filtering the raw data between 0.2-115 Hz using a fourth-order Butterworth filter to remove slow drift, high-frequency noise, and 60 Hz line noise (additionally using a 55-65 Hz stop-band). Broadband high-frequency activity was generated by first calculating the power spectrum density (PSD) from 40-100 Hz (60 Hz line noise removed) with a bin size of 2 Hz and time-step size of 10 ms was estimated using a Hann multi-taper power spectrum analysis in the FieldTrip toolbox (Oostenveld et al., 2011). For each channel, the neural activity between 50-300 ms prior to stimulus onset was used as baseline, and the PSD at each frequency z-scored based on the mean and variance of baseline activity. Single trial broadband high-frequency activity was calculated as the PSD z-scored against pre-stimulus baseline averaged from 40-100 Hz in each 10 ms time step for each trial. Both the single trial potentials (stP) and single trial broadband high-frequency activity (stBHA) were used in all analyses.

Trials with a maximum amplitude five standard deviations above the mean across trials were eliminated, as well as trials with a deflection greater than 25 μV between sampling points. These criteria allow the rejection of sampling error or interictal events, and resulted in elimination of less than 1% of trials when applied in this and previous work (Li et al., 2019).

### Electrode localization

To accurately identify electrode contact location, the co-registration of grid electrodes and electrode strips with cortex was adapted from Hermes et al. (2017). Electrode contacts were segmented from high-resolution post-operative computerized tomography (CT) scans of patients and co-registered with anatomical MRI scans that were conducted before neurosurgery and electrode implantation. This method of using FreeSurfer (https://surfer.nmr.mgh.harvard.edu/, 1999) software reconstructions to co-register with the CT scans accounted for shifts in specific electrode location caused by potential deformation of the cortex (“brain shift” due to cortical displacement by the grid electrode substrate) and resulting signal as a result of grid electrode implantation. Stereotaxic depth electrodes were localized with Brainstorm software (Tadel et al., 2011) that co-registers post-operative MRI with pre-operative MRI images. Complete localization (incorporating the following electrode selection step) is depicted in Figure 1A. The presence of numerically greater HFS contacts in the left hemisphere than right hemisphere is most likely explained by the larger absolute numbers of left than right hemisphere electrode contacts, a result of electrode placement being guided solely by clinical needs of each patient.

### Electrode selection

Electrodes were selected according to anatomical and two functional criteria. Anatomically, electrodes of interest were selected from within ventral temporal cortex below the middle temporal gyrus. Specifically, the midline of the middle temporal gyrus was defined as the upper limit for anatomical consideration: the beginning of the middle temporal gyrus was used to define a posterior threshold, and the midline of the middle temporal gyrus terminating at the temporal pole was used as the anterior threshold for electrode selection. We conducted multivariate classification over data from the localizer session to identify face and word sensitive electrodes (described in next section). Functionally, highly category selective electrodes of interest demonstrated a peak six-way face classification *d’* score greater than 0.8, corresponding to *p* < .01 and a large effect size (Cohen, 1988) for the preferred category (face or word) using a Naïve Bayes classifier (note that in Figure 3 we also examine the robustness of effects to varying thresholds). Electrodes were not considered highly selective if a *d’* score greater than 0.8 resulted from face or word stBHA values with systematically less deviation from baseline relative to other conditions (whereby above chance classification could occur simply by systematically lower response magnitude), resulting in the removal of 7 electrodes across 5 patients. Selective electrodes were also required to show a maximal stP or stBHA response to either faces or words for at least 50 ms during the stimulus presentation period. Electrodes that met these criteria were labeled as highly face selective (HFS) or highly word selective (HWS). Within each patient’s montage, all VTC electrodes of interest that did not meet the criteria for high selectivity for faces were labeled as non-highly face selective (NFS) and those that did not meet this criteria for words were labeled as non-highly word selective (NWS; note that HFS electrodes could be considered NWS and HWS electrodes could be considered NFS).

Finally, to control for any systematic differences in anatomic location between high selectivity and non-highly selective contacts, the most anterior non-highly selective contacts from each montage (which were more numerous and more anteriorly located, by an average 16.4 millimeters, than the selective contacts) were removed until the high selectivity and non-highly selective contacts from each montage were matched anatomically along the anterior-posterior axis. In total, 382 non-highly selective contacts were removed from the 17 patient electrode montages, or 22.47 contacts per montage on average. Before trimming and balancing, non-highly selective contacts were on average 16.4 millimeters more anterior than high selectivity contacts (y = −0.0343 mean placement for non-highly selective compared to y = −0.0507 mean placement for high selectivity contacts). Functionally, this trimming procedure yielded high selectivity and non-highly selective contact populations in each patient’s montage with equivalent mean coordinate values along the anterior-posterior axis and ensured that any latency differences between populations could not immediately be attributed to any expected conduction delays. Indeed, recent work has demonstrated a relationship between response onset latency and situation along the anterior-posterior axis, such that more anterior contacts emerged later in time (Schrouff et al., 2020). Note that this anatomical balancing procedure did not meaningfully alter the time course of classification over non-highly selective contacts compared to retaining all anterior non-highly selective contacts and all results remained similar if non-balanced electrodes were used in the analyses (Supplemental Figure 2). See figure 1A for all electrodes used in further analyses.

Note that the locations of face and word selective electrodes are more distributed than is typically reported in group-level neuroimaging studies (Kanwisher & Yovel, 2006), though they are consistent with the individual variability seen in other imaging modalities (Glezer & Riesenhuber, 2013; Weiner & Grill-Spector, 2013; Gao, Gentile & Rossion, 2018; Zhen et al., 2015; Rossion et al., 2012; Cohen et al., 2002; Dehaene et al., 2004; White et al, 2019) and are consistent with prior iEEG studies (Li et al., 2020, Allison et al., 1999, Hagen et al., 2020; Matsuo et al., 2015; Jacques et al., 2020; Lochy et al., 2018). See Boring et al. (2021) and Figure 1 of Li et al. (2019) for a more thorough examination of the iEEG-derived map of VTC category selectivity, including illustrations of individual subjects from these localizer results and a map that includes all categories used.

### Multivariate classification: Naïve Bayes classifier

We first used a Naïve Bayes classifier with 3-fold cross validation to examine category selectivity over time at individual electrode contacts throughout ventral temporal cortex. Both stP and stBHA signal values were used as input features in the classifier with a sliding 100 ms time window (10 ms width) as previous studies have shown increased sensitivity and specificity when using both stP and stBHA (Miller et al., 2016). Indeed, stP and stBHA metrics have been shown to capture separate and complementary aspects of the physiology that contribute to visual processing as measured with iEEG (Lescynski et al., 2019). stP signal was sampled at 1000 Hz and stBHA at 100 Hz, which yielded 110 features (100 mean stP voltage potentials and 10 normalized mean stBHA PSD values). Thus at each time point at each electrode for each of 3 cross validation steps, the classifier was trained on the first 2-folds and performance evaluated on the left out fold for 6-way classification of the six object categories presented in the localizer session. Trials were divided into three folds by random assignment. We used the sensitivity index *d’* for face or word category against all other categories to determine face and word selective contacts. *d’* was calculated as Z(true positive rate) – Z(false positive rate), where Z is the inverse of the Gaussian cumulative distribution function.

### Elastic net regularized logistic regression

To examine the temporal dynamics of face and word individuation, we used elastic net regularized logistic regression with three-fold cross validation implemented with the GLMNET package in Matlab. Elastic net was chosen as a means to identify diagnostic electrode contacts by removing non-informative and/or highly correlated classifier features. These series of classification problems were conducted iteratively in four different electrode populations: individual face classification from experiment 1 data in VTC HFS contacts and VTC NFS contacts, and individual word classification from experiment 2 data in VTC HWS contacts and VTC NWS contacts (as defined above). Face identity classification was conducted across expression and gaze direction, effectively varying the low-level visual features of each face identity such that this classification problem was not simply face image classification.

stP signal was first downsampled to 100 Hz to yield an equal number of stP and stBHA features. stBHA signal was then normalized with a Box-Cox transformation to ensure that both stBHA and stP were both normally distributed. Thus at each time point, stP and stBHA values from each trial were arranged as a *P*-dimensional vector corresponding to 2 * number of contacts in each of the four predefined electrode contact populations. The time course of face and word individuation was identified by examining the pairwise decoding accuracy of a classifier using 3-fold cross-validation. The regularization parameter (α) was set a priori to 0.9 (range of 0-1) to favor more sparse classification solutions that produce more statistically interpretable results (similar to applying a lasso (L1) penalty in the case of α = 1) while avoiding degeneracies that sometimes emerge in full L1 regularization (Friedman et al, 2008). The results of this analysis are depicted in Figure 2. For display purposes, group mean time courses were smoothed with a moving average of 30 millisecond fixed window length.

For comparison purposes, L1 regularized logistic regression (α=1) was also repeated in the same manner as the above elastic net analyses (classification conducted separately for highly and non-highly category selective populations) to demonstrate minimal difference in the time course of *d’* values from the different regularization procedures.

To demonstrate the robustness of general trends of individuation to the selection criteria for highly and non-highly selective contact populations, the elastic net classification procedure was repeated with additional thresholds determined by dividing face and word contact populations into partitions of equal numbers. To do so, all contacts across all subjects in face and word tasks, respectively, were sorted according to peak d’ selectivity value from the category localizer. Then, these contacts were divided into six equal partitions. Then, elastic net regularized classification was conducted again according to the following groupings: 1) bottom two partitions labeled as NFS, top four partitions labeled as HFS (corresponding d’ value of 0.61 dividing the two groups); 2) bottom three partitions labeled as NFS, top three partitions labeled as HFS (corresponding d’ value of 0.7 dividing the two groups); 3) bottom four partitions labeled as NFS, top two partitions labeled as HFS (corresponding d’ value of 0.82 dividing the two groups). This procedure was repeated for word selective contacts at the following d’ thresholds: 0.58, 0.67, 0.86. The results of this analysis are depicted in Figure 3. For display purposes, group mean time courses were smoothed with a moving average of 30 millisecond fixed window length. The partitions corresponding to the bottom 1/6 and top 5/6 (and vice versa) are not demonstrated because not all subjects had contacts in the lowest and highest partitions.

### L1 regularized logistic regression

To examine the diagnosticity of brain activity from highly and non-highly category selective electrode populations in concert with one another, we repeated the above classification analyses with L1 as opposed to elastic net regularization and examined the proportion of electrode contacts that were entirely penalized and removed from the classifier model. Additionally, all VTC electrode contacts (highly and non-highly category selective) were used to train each classifier, as opposed to splitting the electrode populations as in the previous analyses. After conducting pairwise face classification and pairwise word classification, the classifier weights from each pairwise classification for each electrode contact were extracted and the number of non-zero (positive or negative) weights for each contact tabulated. The percent of electrode contacts with non-zero weights was determined at every time point after baseline normalization.

Baseline normalization consisted of determining the threshold of non-zero weight counts that would yield <1% contacts with non-zero weights during the baseline period. The total percentage of electrode contacts assigned non-zero weights for at least 50 ms across the entire time course was determined, and results from this analysis are depicted in Figure 4A. This change in classifier does not alter the time course of individuation compared to the original elastic net procedure.

### Electrode Diagnosticity in non-highly category selective areas

Having examined the contributions of highly and non-highly face and word selective contacts to exemplar representation, we were then interested in examining whether 1) non-highly face and word selective sites with selectivity for a different category differed in their contributions to exemplar representation from 2) non-highly face and word selective sites lacking any other category selectivity. The main question here is the extent to which contacts that demonstrate category selectivity will contribute to exemplar representation for a different category. Thus in addition to examining highly and non-highly category selective contacts, we further decomposed the non-highly face and word category selective populations into two sub groups: other category selective (OCS) and not significantly selective for any category (NCS). Face OCS contacts, while not showing high selectivity for face images, did show selectivity for either word, house, body, or hammer stimuli based on the same criteria for selectivity described above. Word OCS contacts did not show high selectivity for words, but did show high selectivity for either face, house, body, or hammer stimuli. NCS contacts showed no high selectivity for any of the categories presented in the localizer task. Category selectivity for non-face and word categories was established with the same method of Multivariate Naïve Bayes classification at the category level as previously outlined, and weights extracted in the same method outlined in the immediately preceding section.

To further quantify unique diagnosticity in highly and non-highly selective populations and address concerns of volume conduction or signal bleed from high selectivity populations, multivariate linear regression was carried out to regress the high selectivity signal from non-highly selective signal in each subject’s dataset. Following this, the same elastic net regularized logistic regression procedure was used to classify individual stimuli across time using the residuals of the non-highly selective signal.

Where the above analyses examine whether unique information is present in non-highly selective regions, a representational similarity analysis (RSA) was performed to examine the overlap in information between highly and non-highly selective populations. Confusion matrices from pairwise classification accuracies as measured with elastic net regularized logistic regression between highly and non-highly selective populations for faces and words were first calculated. For each subject, a vector corresponding to the lower matrix diagonal from pairwise classification accuracies was extracted separately for highly and non-highly selective populations and a Spearman correlation computed between them. This correlation measures whether pairs of faces or words that were easy or difficult to classify from one another in high selectivity contacts were also easy or difficult to classify from one another in non-highly selective contacts.

## Statistical Analyses

For the category localizer with Naïve Bayes classification, row permutation tests on a subject level were used to establish a *d’* threshold for category selective contacts. For each subject within each permutation, the condition labels for each trial were randomly shuffled and the same classification procedure as above was used 1000 times for a randomly selected channel in each electrode montage. The peak *d’* value from each permutation was aggregated into a group-level distribution comprising the null distribution from each permutation for each subject. The *d’* value corresponding to *p* < .01 was estimated from this histogram and used as a selectivity threshold to determine highly and non-highly selective contact populations for each subject.

For face and word individuation as measured with elastic net regularized logistic regression, row permutation tests were used to establish a significance threshold for classification accuracy for each subject. For each permutation, a classifier model was optimized and test condition labels shuffled to test model predictions on randomized data. This procedure was repeated 1000 times to generate a null distribution. The true classification values and null distributions for each subject were combined into group-level distributions, and the mean true classification value and mean null distribution compared to one another. Classification accuracy was deemed significant at a level of *p* < .05 with FDR correction (Benjamini-Hochberg procedure for dependent tests), with a minimum temporal threshold of 3 contiguous significant time points. Thus, although different subjects contributed different numbers of contacts to each classification analysis, all subjects are weighted equally in the group mean depicted in Figure 2.

Onset sensitivity was determined by with 3 metrics examining the individual subject-level statistics. For the first method, the same true classification values and null distributions from above were compared on an individual level, and the first time point significant at a level of *p* < .05 with FDR correction (Benjamini-Hochberg procedure for dependent tests) with a minimum temporal threshold of 3 contiguous significant time points was used as the onset marker for each subject. Vectors of onset markers compiled from all subjects were compared between HFS / NFS, and HWS / NWS electrode populations with paired-sample t-tests. Because this method is somewhat sensitive to the magnitude of the response (e.g. higher magnitude will cross the statistical threshold sooner) two other methods for calculating onset that are more robust to magnitude differences were used as well.

The second onset determination method was adapted from Schrouff et al. (2020): for each subject, the time course of mean classification values for each classification problem (HFS, NFS, HWS, and NWS) were normalized to peak classification value, and a sliding window with 50 ms bins and 10 ms overlap was implemented. Classification average and standard deviation in the baseline period of −100 to 0 ms was estimated, and the first period with 3 contiguous bins surpassing the baseline threshold was marked as the signal onset for a given subject’s classification time course. Vectors of onset markers compiled from all subjects were compared between HFS / NFS, and HWS / NWS electrode populations with paired-sample t-tests. Schrouff et al (2020) show that this method for finding onset times is robust to differences in peak magnitude across comparisons.

For the third onset determination method, onset sensitivity was measured as the first 3 contiguous time points where classification values for each subject were greater than 25% of the peak value. Vectors of onset markers compiled from all subjects were compared between HFS / NFS, and HWS / NWS electrode populations with paired-sample t-tests. While 25% of the peak value is not necessarily a strict measure of “onset,” it is independent of peak magnitude and provides a metric of whether any differences in peak time are due to differences in slope or whether there is differences in onset (e.g. earlier peak times could be due to sharper rising slope or earlier onset).

To assess significance for the residual classification procedure, the same row permutation test from elastic net regularized logistic regression significance testing above was again used in this context. For the pairwise classification accuracy RSA, a two-sided t-test was used to identify Spearman’s rho values significantly greater than 0.

## Acknowledgements

We would like to thank the patients, their families, and nurses, staff, and physicians at the Epilepsy Monitoring Unit and the University of Pittsburgh Comprehensive Epilepsy Center at the University of Pittsburgh Medical Center, without whom this study would not be possible. We would also like to thank Michael Ward, Sean Walls, and Ellyanna Kessler for assistance in data collection and Julie Fiez for assistance with design of the word experiment. Additional thanks to Chris Baker, Brad Mahon, and Alex Martin for critical comments and feedback on this work. This work was supported by the National Institutes of Health (R01MH107797 and R21EY030297 to A.G) and the National Science Foundation (Graduate Research fellowship to B.B.B., 1734907 to A.G.).

## Competing Interests

The authors declare no competing interests.

## Author Contributions

A.S.G designed the experiment. R.M.R. conducted surgical implantations. B.B.B., M.J.B., R.M.R., and A.S.G. collected experimental data. B.B.B. and M.J.B. analyzed the data and generated figures. B.B.B. and A.S.G. wrote the manuscript.

## Data and materials availability

All data and code is available upon reasonable request to A.S.G.

---

**Supplementary Fig. 1.**
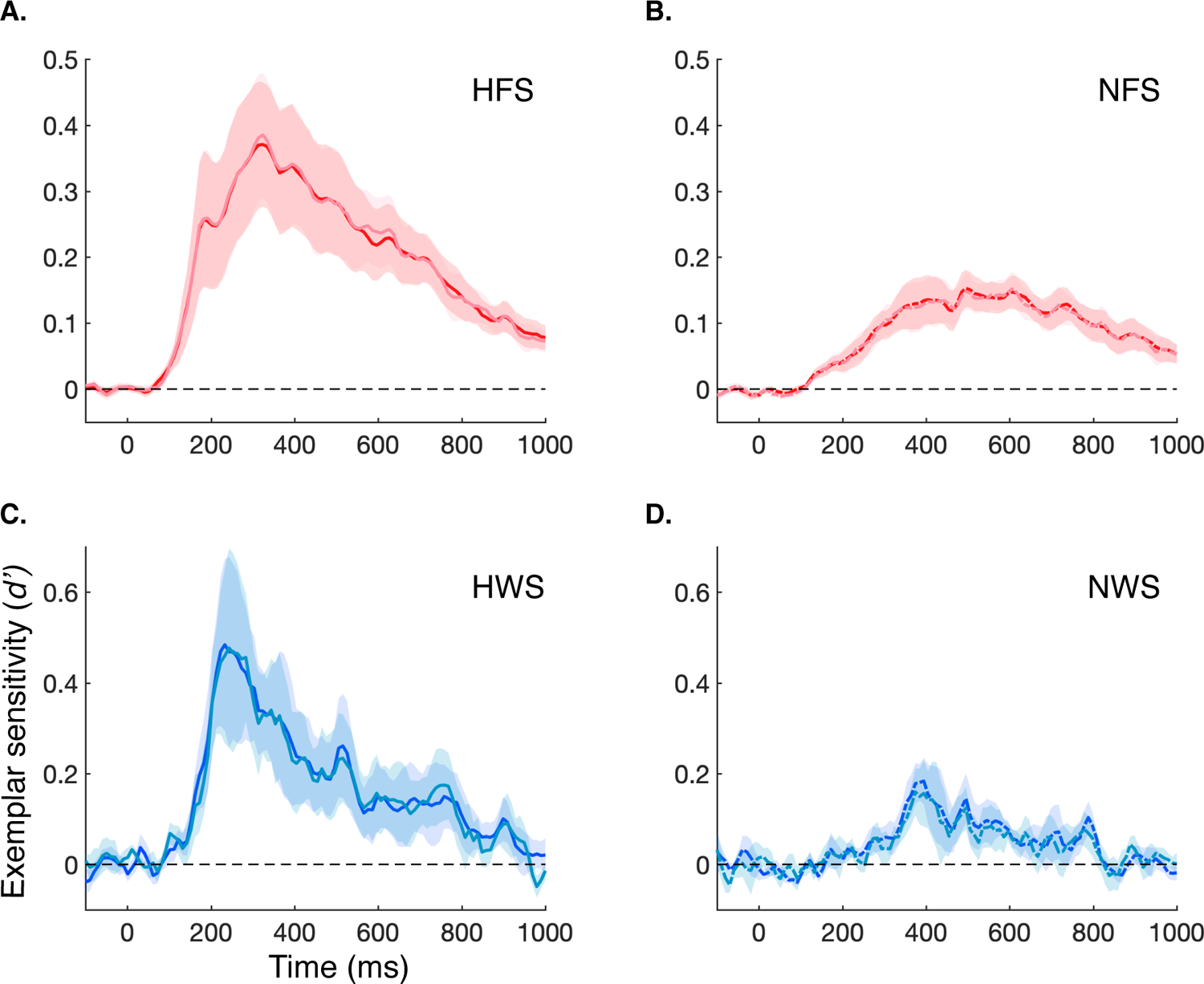
Time course of *d’* exemplar sensitivity as measured with elastic net vs. L1 regularized logistic regression. A) HFS contacts measured with elastic net (dark red) versus L1 (light red) penalization. B) NFS contacts measured with elastic net (dark red) versus L1 (light red) penalization. C) HWS contacts measured with elastic net (dark blue) versus L1 penalization (light blue). D) NWS contacts measured with elastic net (dark blue) versus L1 (light blue) penalization.

**Supplementary Fig. 2.**
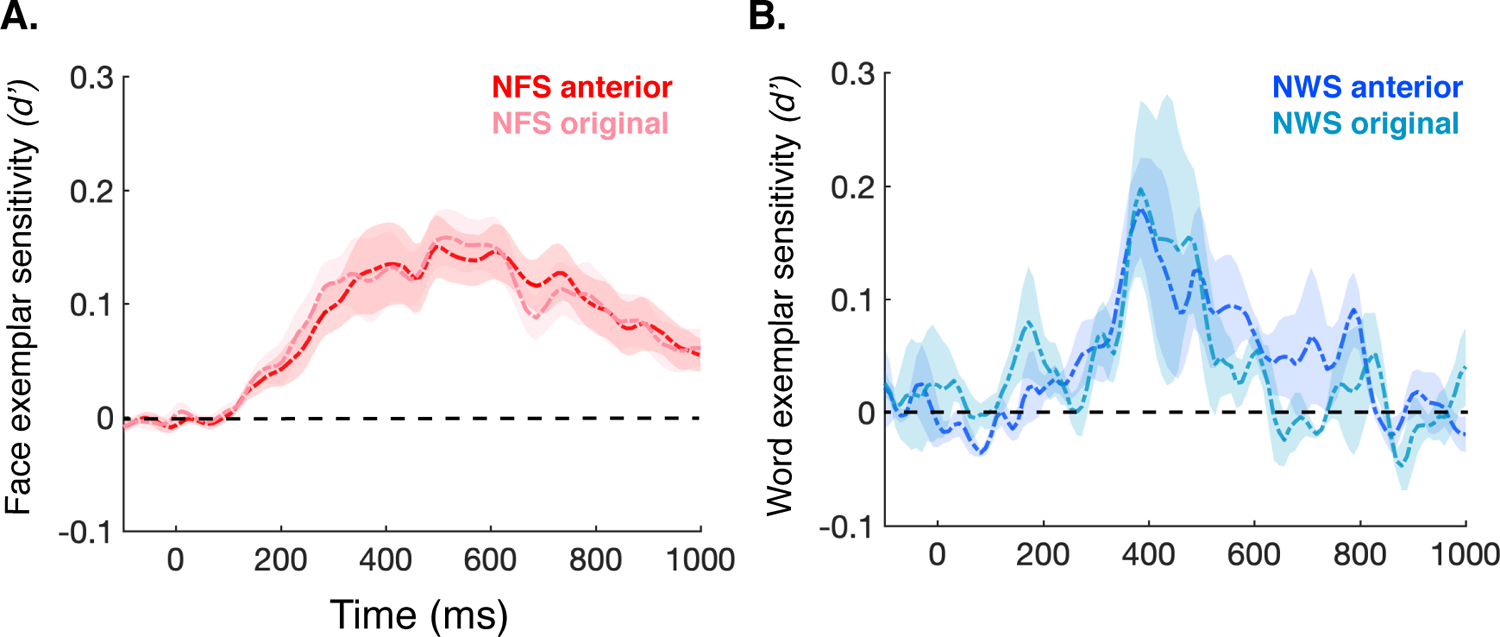
Time course of exemplar sensitivity in original and anterior-inclusive NFS/NWS populations. The same elastic net regularized logistic regression analysis was repeated over the anterior-inclusive contacts for face and word classification. No statistically significant differences in the onset time, peak time, or peak *d’* between original and anterior-inclusive contacts emerged. A) Mean *d’* for NFS original contacts (light red) that peaked at 520 ms (*d’* = 0.17) compared to the mean *d’* for NFS anterior-inclusive contacts (dark red) that peaked at 495 ms (*d’* = 0.15). B) Mean *d‘* for NWS original contacts (light blue) peaked at 395 ms (*d’* = 0.2) after stimulus onset, temporally identical to the mean *d’* for NWS anterior-inclusive contacts that peaked at 395 ms (*d’* = 0.18).

## Notes

### Competing Interest Statement

The authors have declared no competing interest.

### Summary of Updates

additional stP / stBHA recording figures, supplemental analyses, interpretation of findings

